# Challenges in inferring breathing rhythms from olfactory bulb local field potentials

**DOI:** 10.1101/2024.11.08.622727

**Authors:** Sidney Rafilson, Nathan Hess, Teresa M. Findley, Matthew C. Smear

## Abstract

Odors convey useful navigational and episodic information, yet much of the chemical world remains inaccessible without active sampling through sniffing. Respiratory cycles control odor dynamics within the nose, so understanding olfactory bulb (OB) neural dynamics requires accurate respiratory measurements. While respiratory behavior can be measured directly with a variety of chronic methods, these methods are invasive and none are perfectly robust. OB local field potentials (LFPs) have long been known to couple with respiration. Here we investigated whether the precise timing and frequency of respiration can be inferred from OB LFPs. Our results replicate previous findings that OB LFPs across multiple frequency bands align with respiratory cycles. Further, these OB rhythms are locked to time in the respiratory cycle, and not phase. In addition, we show that 2-12 Hz LFP oscillations effectively track sniffing rate. However, a monotonic relationship between LFP-respiratory delay and sniffing rate, which varies across animals, renders the recovery of precise respiratory events challenging. This work underscores the complex and individualized relationship between rodent respiration and OB LFPs, contributing to our understanding of how respiration controls olfaction.

## INTRODUCTION

Sensation in the rodent olfactory system is modulated by respiratory behavior throughout the brain (Wachowiak, 2011). Beginning with inhalation, which drives the binding of odorants with olfactory sensory neurons (OSNs; Kleene, 2008), the respiratory cycle further organizes downstream spiking activity. Sniffing modulates the temporal dynamics of the spatial map of odor identity at olfactory sensory inputs, the glomerular layer (Spors et al., 2006; Verhagen et al., 2007). In addition to this spatial organization, mitral and tufted (M/T) cells are tuned to timing in the respiratory cycle, such that odor representations are temporally structured by sniffing (Cury & Uchida, 2010; Macrides & Chorover, 1972; Shusterman et al., 2011); and this temporal structure is modulated by odor identity (Iwata et al., 2017) and behavioral state (Sterrett et al., 2025). Optogenetic experiments have established that the olfactory system can read out the timing of OSN activity (Chong et al., 2020; Haddad et al., 2013; Li et al., 2014; Smear et al., 2011, 2013). Because of the respiratory alignment of OB population dynamics, analyzing sniff-centric spiking activity has become a powerful tool for understanding the encoding of odor identity and intensity (Cury & Uchida, 2010; Wilson et al., 2017), as well as the effects of behavioral state and context on sensation (Sterrett et al., 2025).

Concurrent with the fast activity of single unit firing are slower voltage fluctuations known as local field potentials (LFPs). LFPs represent the sum of sub-threshold electrical activity of hundreds to thousands of cells (Buzsáki, 2019) and are thought to be a measure of collective neuronal dynamics, particularly synaptic activity (Buzsáki et al., 2012). Despite measuring a fundamentally different aspect of neuronal activity, OB LFPs are coupled with respiratory behavior across multiple frequency bands (Rojas-Líbano et al., 2014; Rojas-Líbano & Kay, 2008) and relate dynamically to odor learning, sensitization, and discrimination (Kay et al., 2009). As such, accurately measuring respiration to analyze LFPs within the context of a sniff is essential for understanding the collective neuronal activity in the OB and describing the degree of alignment between respiration and OB LFPs.

There are numerous methods of measuring respiration in rodents. All of these techniques require additional surgeries and hardware to be added to already-challenging chronic electrode array implantation. Further, each has its unique drawbacks. Pressure sensor-coupled chambers for plethysmography provide precise measures of respiration but require a small recording chamber, restricting animal mobility (Lim et al., 2014). Intranasal pressure or temperature sensors can solve the problem of animal mobility while maintaining high signal fidelity (Casali et al., 2024; Grimaud & Murthy, 2018); however, implanted sensors are surgically invasive, particularly around the OSNs, and prone to blockage (McAfee et al., 2016). of the diaphragm measures respiration without the drawback of blockage or injury to the nasal epithelium, but are extremely invasive (Grimaud & Murthy, 2018). To address the issues of invasiveness and mobility while maintaining high signal fidelity, a method involving a chronically implanted temperature sensor, or thermistor, was devised. The thermistor is implanted into the hollow space above the anterior portion of the nasal cavity, safely avoiding the epithelium and nasal passages while still capturing respiration-driven temperature fluctuations (Findley et al., 2021; McAfee et al., 2016). While the thermistor provides a high-fidelity recording in mobile animals and minimizes invasiveness, the method still requires a craniotomy, and it suffers from signal degradation in recordings spanning weeks or months. Even at their best, all these methods entail additional hardware and wires to be added to the preparation.

To circumvent these complexities and motivated by the close alignment between OB LFPs and respiration (Kay et al., 2009; Rojas-Líbano et al., 2014; Rojas-Líbano & Kay, 2008), we asked whether respiration can be inferred from OB LFPs, making direct respiratory measurements unnecessary. To use LFP as a proxy for respiration, we sought to determine whether a model could translate LFPs into sniffing parameters, such as inhalation onset and sniff rate. In doing so, sniff-centric neural data may be analyzed without the need for direct respiratory measurement, but rather through a respiratory reconstruction from OB LFPs.

Our results replicate previous findings that theta band (2–12 Hz) LFP oscillations track respiratory rhythms and that their power spectra track instantaneous sniffing rate (Rojas-Líbano et al., 2014). In the time domain, inhalation-aligned theta and gamma (65 – 100Hz) oscillations exhibited strong correlations with respiratory cycle length. However, considerable inter-mouse and intra-session variability in the latency between inhalation onset and LFP peaks hindered the accurate recovery of inhalation timing. While we found that time-domain methods outperformed frequency-domain approaches in estimating sniffing rate, especially in freely moving conditions, precise cycle-by-cycle reconstruction remains limited. These findings emphasize the potential and limitations of using OB LFPs as a proxy for respiration and motivate future work to understand the complex coupling of neural dynamics and active sampling.

## METHODS

### Animal housing and care

All procedures were conducted in accordance with the ethical guidelines of the National Institutes of Health and were approved by the Institutional Animal Care and Use Committee at the University of Oregon. Animals were maintained on a reverse 12/12 h light/dark cycle. All recordings were performed during the dark phase of the cycle. Mice were exposed to ambient light during experimentation. Mice were C57Bl6/J background and were 8–12 weeks of age at the time of surgery.

### Surgical procedures

Animals were anesthetized with isoflurane (3% concentration initially, altered during surgery depending on response of the animal to anesthesia). Incision sites were numbed prior to incision with 20 mg/mL lidocaine. Thermistors were implanted between the nasal bone and inner nasal epithelium (Findley et al., 2021). A custom titanium head bar and Janelia micro drive were implanted.

For olfactory bulb array implantation, we administered atropine (0.03 mg/kg) preoperatively to reduce inflammation and respiratory irregularities. Surgical anesthesia was induced and maintained with isoflurane (1.25–2.0%). Skin overlying the skull between the lambdoid and frontonasal sutures was removed. A rectangular window was cut through the skull overlying the lateral half of the left bulb for insertion of the recording array. The array was lowered to a depth of 1mm and cemented in place with Grip Cement. To minimize postoperative discomfort, Carpofen (10 mg/kg) was administered 45 minutes prior to the end of surgery. Mice were housed individually after the surgery and allowed 7 days of post-operative recovery.

### Behavioral recordings

Mice were restrained by head fixation then placed in a 15 cm by 40cm behavioral arena. After a period of head-fixation, mice were released and permitted to freely move around the arena, without explicit training or reward structure, while breathing and neural data was collected. We record sniffing using intranasally implanted thermistors (TE Sensor Solutions, #GAG22K7MCD419), amplified initially with custom-built op amp (Texas Instruments, #TLV2460, circuit available upon request) and then a CYGNAS, FLA 01 amplifier fed into the analog input of an OpenEphys box.

#### Electrophysiology

Following 3-day recovery period post-surgery mice were head fixed and the custom microdrive was advanced to the regions of interest (ROI) while recording. Either Si probes (Diagnostic Biochip P-64-7) or a custom implanted array of 8 tetrodes passed in pairs through 4 linearly-aligned 27-gauge stainless steel hypodermic tubes. Tetrodes were made of 18 µm (25 µm coated) tungsten wire (California Fine Wire). Once the ROI was reached a minimum of 24 hours was allowed prior to data collection to increase recording stability. Data were acquired via a 128-channel data acquisition system (RHD2000; Intan Technologies) at a 30 kHz sampling frequency and OpenEphys software (http://open-ephys.org). Custom Bonsai code was used to align the respiratory and the electrophysiology recording captured with no filters applied in the OpenEphys software.

## DATA ANALYSIS

All data was analyzed in custom python scripts (Smear-Lab/sniff-LFP: Local Field Potential Project-- Smear Lab). LFP and thermistor preprocessing for all analysis consisted of lowpass filtering the signal below 300Hz using an order 4 Butterworth filter, followed by down sampling to 1 kHz, using the SciPy signal processing library. Inhalation times were extracted by finding peaks in the temperature signal after smoothing with a 25ms moving Savitsky-Golay filter. Sniffs with a duration less than the 5th and greater than the 95th percentile were excluded from the analysis.

### Spectral Analysis

Power spectrum and spectral coherence were estimated between individual channels of the LFP and the thermistor signal. To build time-frequency spectrograms, we used 50% overlapping 4s windows. This choice of window lengths yielded a lower bound on frequency content at 0.25 Hz and given the sampling rate, a Nyquist frequency of 500Hz. To home in on the frequency band of interest, we bandpass filtered the LFPs using another order 4 Butterworth filter. To limit spectral leakage in the short windows, the power spectrum estimates were computed using Thompsons Multitaper procedure (Thomson, 1982). This method involves defining a time-bandwidth product *NW* where *N* is the window length and *W* is the half-bandwidth of interest. Then, *K* orthogonal data tapers from the Slepian sequence (Slepian, 1978) are computed for each window. Typically, *K* ≈ 2*NW* – 1 and hence we chose to use 7 tapers. Finally, averaging the Fourier transforms of the time series by each of these tapers gives the final spectral estimate for the *n*^th^ window:

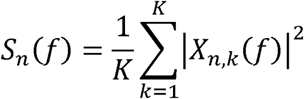

Where *X_n,k_*(*f*) is the Fourier transform for the *n*^th^ window and *k*^th^ taper such that

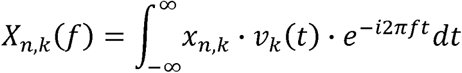

With *X_n_*_,*t*_ (*t*) the signal in the *n*^th^ window and *v_k_* (*t*) the *k*^th^ data taper. Spectral coherence was calculated as follows:

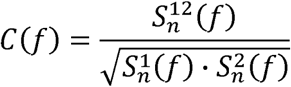

Where 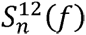 corresponds to the estimated cross-spectrum calculated by multiplying the spectral estimate of the first signal 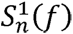 with the complex conjugate of the spectral estimate of the second signal 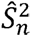.

Time-frequency heatmaps represent coherence or spectral power by color with a linear color mapping fit to the middle 90% of the data range. Confidence limits on coherence were created by circularly shifting the thermistor signal before computing the windowed coherence. At least 1000 pseudo-random circular shifts were performed and a z-score threshold of 1.65 from the null coherence distribution was used as a lower limit on significant coherence.

### Time-domain Analysis

In the time-domain we analyzed wavelengths and latencies between inhalation times and peaks in the LFPs. To reliably detect peaks, we averaged unit variance scaled LFP epochs aligned to inhalation times, within sniff rate bins. The bins were linearly spaced between 2 and 12Hz with a width of 0.5Hz, thus discretizing the sniff rates into 20 partitions. We detected peaks in the averaged LFP epochs using the SciPy find_peaks function with prominence of 0.1 and a minimum distance set to the reciprocal of the upper passband. This method allowed us to estimate the distributions of mean inhalation-to-peak latencies and LFP wavelengths across sniff rates, mice, and sessions.

### Respiratory frequency inference

We inferred the average sniff rate in sliding windows ranging from 1 to 128 seconds using time and frequency domain approaches. In both methods, we worked with the 2 – 12Hz bandpass filtered LFPs. In the time domain we took the mean reciprocal of the inter-peak-intervals. We infer the frequency of the *w*^th^ window in the time domain by,

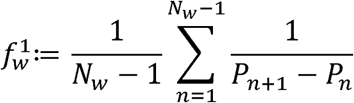

*Where P_n_* is the time of the *n*^th^ peak and *N_w_* the number of peaks in the associated window. The frequency domain method finds the frequency component of the power spectrum between 2 and 12 Hz with the maximum power. That is,

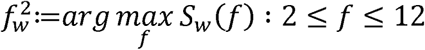

Where *S_w_* is the power spectrum in the associated window.

## RESULTS

Given the importance of respiration and the challenges of capturing it directly, we attempted to reconstruct respiration from OB LFPs in the frequency and time domains. We first extracted the respiratory signal and LFPs in three frequency bands (Figure 1A, 1B). Time-frequency spectrograms of the unfiltered LFPs show that the peak in the power spectrum tracks instantaneous sniffing rate, as defined by the reciprocal of inter-peak-intervals in the thermistor signal (Figure 1C *top*). Likewise, instantaneous sniff rate tracks the maximum power in the thermistor power spectrum (Figure 1C *middle*). Taking the linear correlation between the raw LFPs and thermistor power spectra demonstrates significant spectral coherence between OB theta oscillations and respiration (Figure 1E). Further, the coherence tracks instantaneous sniffing rate (Figure 1C *bottom*) and reaches significance with respect to a null distribution in a narrow window associated with the instantaneous sniff rate distribution, approximately 2 – 12Hz (p < 0.05, permutation test; Figure 1D, 1E). These results reproduce previous studies showing that OB theta oscillations track breathing rhythms (Rojas-Líbano et al., 2014).

**Figure 1:**
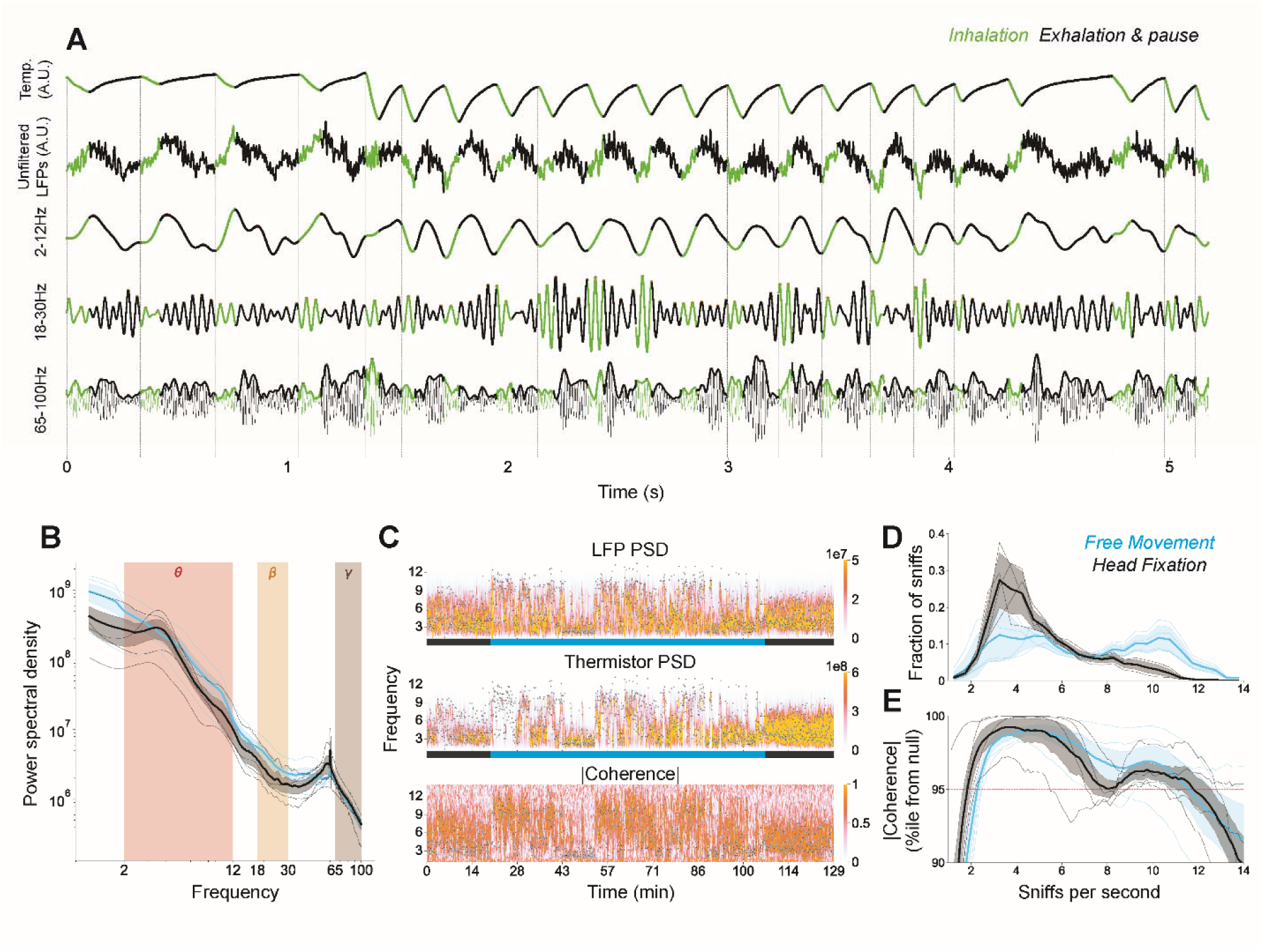
Spectral analysis reveals coherence between LFPs and sniffing. **A)** Sniffing and neuronal signals ***Top)*** Thermistor temperature for recording respiration. ***Second)*** Unfiltered LFPs. ***Third)*** 2 – 12Hz filtered LFPs (Theta). ***Fourth)*** 18-30Hz filtered LFPs (Beta). ***Bottom)*** 65-100Hz filtered LFPs (gamma) and the envelope (gamma envelope). Signals are colored green during inhalation and black during exhalation and pause. **B)** LFP power spectral density. Thin lines represent individual mice with the thicker line to represent the grand means. Shaded region is the standard error of the mean across mouse means. Color shows head fixation in black and free movement in blue. **C)** Time-frequency Spectrograms computed using Thompson’s multitaper method. Black scatter overlay represents instantaneous sniff rate. For visual clarity, only a subset of instantaneous rate points are shown. The color bar shows head fixation in black and free movement in blue. Example session is shown. ***Top***) PSD of raw LFPs. ***Middle***) PSD of thermistor signal. ***Bottom***) Absolute spectral coherence between LFP and thermistor signals. **D)** Instantaneous sniff rate distributions, color scheme as in B. **E)** Absolute coherence represented as a percentile from 1000 circularly shifted null distributions, color scheme as in B. Red dotted line represents the 95% confidence threshold.

To capture the relationship between OB LFPs and respiration in the time and frequency domains, we visualized LFP epochs centered on inhalation onset and examined the sequences of various features across the epochs. First, we observed alignment of unfiltered LFPs to inhalation onset via a constant time delay to the first prominent peak (Figure 2A), whereas aligning to sniff phase revealed a drift in the LFP alignment (Figure 2E). Further, the LFP wavelength between successive prominent peaks appears to be correlated with respiratory cycle length (Figure 2A). Bandpass filtering between 2 and 12Hz shows that the unfiltered LFP-respiratory relationship reflects the theta range (Figure 2B, 2F). In a minority of sessions, we observed beta oscillations (18-30Hz) aligning to inhalation onset by a latency to peak and wavelength invariant of sniff frequency (Figure 2C). Aligning to phase revealed another drifting relationship in the LFP alignment (Figure 2G). Finally, we bandpass filtered the LFPs between 65 and 100 Hz and aligned the amplitude envelope of these gamma oscillations to inhalation onset, finding similar alignment as in the theta band (Figure 2D, 2E). We noticed no differences in latency to peak or LFP wavelength between free movement and head fixation. These visualizations reveal the time-domain alignment of LFPs with respiration: LFPs align to absolute time in the respiratory cycle, and not the relative time as has been proposed for OB spike coding (Shusterman et al., 2011; Smear et al., 2011).

**Figure 2:**
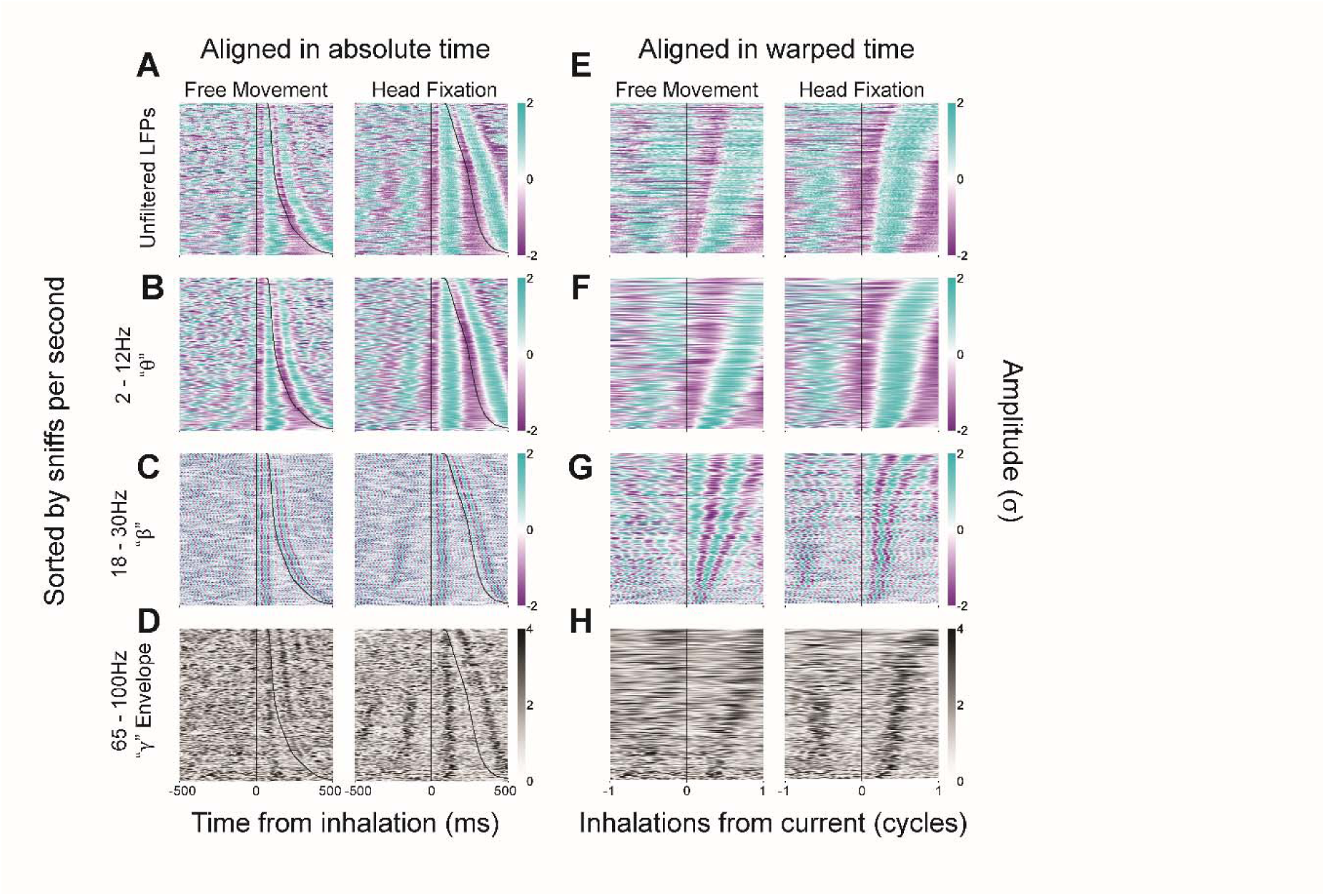
Inhalation aligned LFP visualizations reveal respiratory coupling in absolute time and not phase. Colormaps show inhalation time aligned LFP epochs sorted in the y-direction according to the instantaneous sniff rate from which they are aligned. Color represents the unit variance scaled amplitude. **A, E)** Unfiltered LFPs. **B, F)** 2 – 12Hz filtered LFPs (Theta). **C, G)** 18-30Hz filtered LFPs (Beta). **D, H)** 65-100Hz filtered envelope (gamma envelope). **A, B, C, D** are aligned in absolute time. **E, F, G, H** are aligned in warped time (phase). Example session shown.

To quantify the relationship between LFPs and respiration in the time-domain, we constructed distributions of the delay to the first prominent LFP peak and inter-peak-interval, across the inhalation aligned epochs. We predicted that the primary LFP peak in each respiratory cycle would occur at a reliable latency after inhalation. However, we observed substantial variability in the number of peaks per respiratory cycle across all frequency bands (Figure 1A). Because of this variance, understanding the distributions of delays and cycle lengths was made challenging. While the visualizations appear to show a systematic timing relationship between the LFP epochs and respiration, the variance in the number of prominent peaks per respiratory cycle suggests that this relationship may not be robust on a sniff-by-sniff basis.

To uncover time-domain relationships which may be masked on a sniff-by-sniff basis, we averaged the unit variance scaled LFP epochs aligned to inhalation times, within sniff frequency bins (0.5Hz), and detected peaks on these averaged LFP epochs. In the theta band, 98% and 92% of bins contained a single peak during free movement and head fixation, respectively (Figure 3B). Likewise, the gamma envelope contained single peaks in 93% and 91% of bins (Figure 3D).Using these averaged and normalized epochs, we found that respiratory and LFP cycle lengths were strongly correlated in both theta band (free movement: r = 0.999, p < 0.01; head fixed: r = 0.999, p < 0.01; Figure 3B) and gamma envelope oscillations (free movement: r = 0.968, p < 0.01; head fixed: r = 0.996, p < 0.01; Figure 3D). Similarly, respiratory cycle length and latency to peak showed significant monotonic correlations in both theta band (free movement: ρ = 0.983, p < 0.01; head fixed: ρ = 0.953, p < 0.01; Figure 3B) and gamma envelope (free movement: ρ = 0.994, p < 0.01; free movement: ρ = 0.922, p < 0.01; Figure 3D). While these relationships were consistent at the population level, correlation strengths varied across individual mice in free movement (theta: r = 0.991-0.999; gamma: r = 0.451-0.991) and head fixed conditions (theta: r = 0.992-0.999; gamma: r = 0.909-0.996). In contrast, beta band activity showed no significant correlation with respiratory cycle length in either cycle length (free movement: r = 0.491, p = 0.03; head fixed: r = 0.553, p = 0.02) or peak latency (free movement: ρ = 0.487, p = 0.03; head fixed: ρ = −0.022, p = 0.93; Figure 3C).

**Figure 3:**
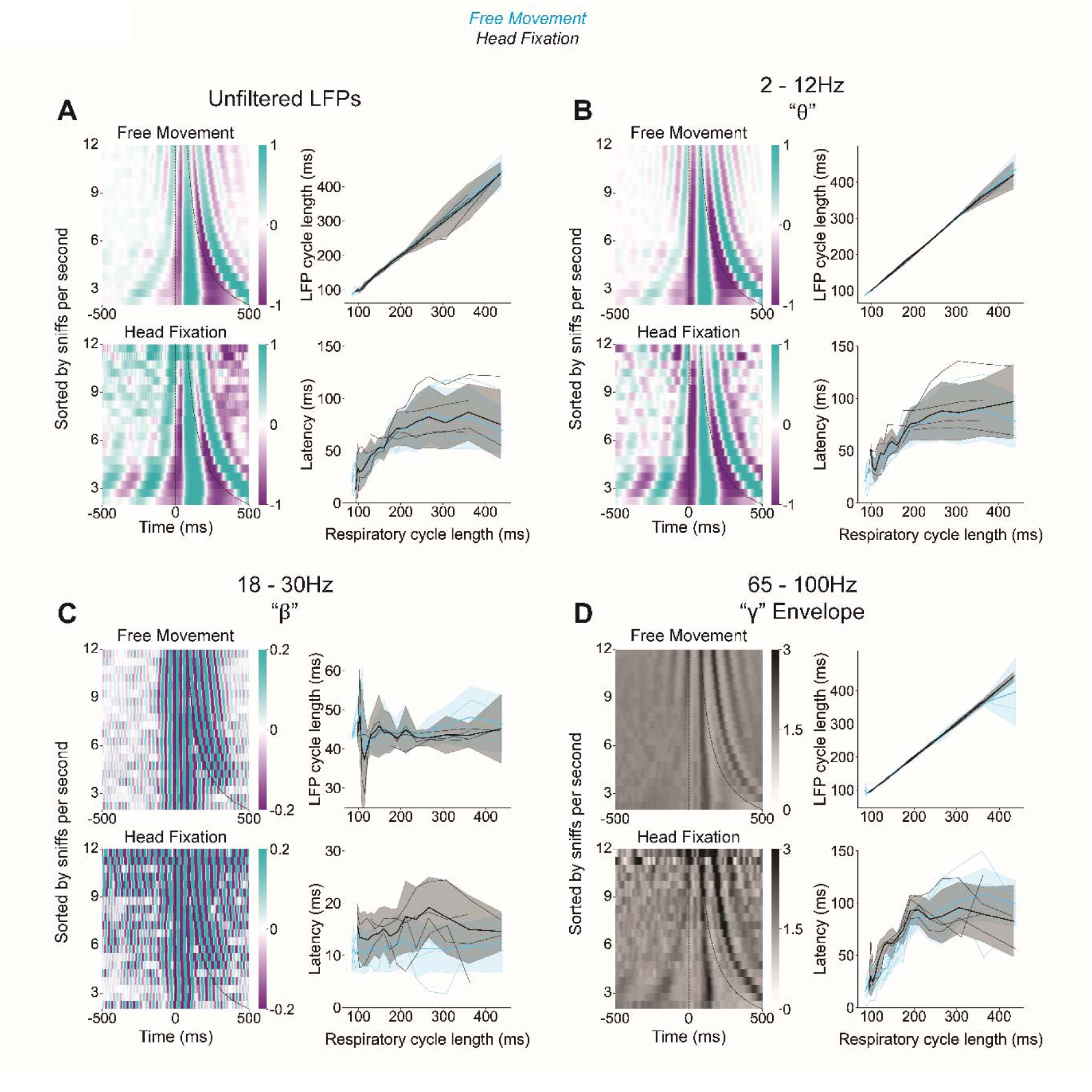
Mean inhalation aligned LFP epochs are correlated with respiration parameters. Colormaps show mean inhalation time aligned LFP epochs in 0.5Hz bins, sorted in the y-direction according to the sniff rate bin from which the epochs are averaged in. Visualizations show one example session. Right column line plots show the LFP cycle length and latency from inhalation time to peak in LFP as a function of respiratory cycle length, across all mice and sessions. **A)** Unfiltered LFP cycle length is linearly correlated with respiration cycle length and the time latency to peak in the LFP has a non-linear monotonic relationship with respiratory cycle length. **B)** 2 – 12Hz filtered LFPs (Theta) rhythms show the same time-domain characteristics as raw LFPs with respect to respiration, but with stronger correlations. **C)** 18-30Hz filtered LFPs (Beta) cycle lengths and latency to peak are invariant to respiratory cycle length. **D)** 65-100Hz filtered envelope (gamma envelope) oscillations have similar time-alignments to respiration as in the theta band, although with a stronger cycle length correlation in head fixed compared to free moving.

Because of the spectral coherence and time-domain relationships between LFPs and sniffing we asked if the precise timing of inhalation and the respiratory frequency could be inferred from the LFPs. The across-mouse variation in the mean latency from inhalation onset to peak in the LFPs renders recovering the precise timing of inhalation onset non-generalizable across mice (Figure 4A, 4B). However, the strong correlation between LFP and respiratory cycle length (Figure 3B) suggests that the respiratory frequency can be recovered from the theta oscillations. To test this hypothesis, we devised time and frequency domain methods of inferring respiratory frequency using sliding windows of LFP. The time-domain method finds the mean inter-peak-interval in the LFP window whereas the frequency domain method estimates the frequency with maximum power. Averaging the results across all mice, sessions and bin sizes we found that the time-domain method outperformed the frequency method (p < 0.001, rank sum test; Figure 5E, 5F), and both models performed better during free movement compared to head fixation (p < 0.001, rank sum test) Notably, during free movement using the time-domain method with 8+ second windows, we estimated sniff rate within 1Hz of the true value in over 75% of windows (Figure 5C). Despite the increasingly accurate estimates with increased window size using the time domain method (Figure 5C, 5D), accuracy in the frequency domain model peaks at 60% in 2 second windows, and longer windows lead to less accurate predictions (Figure 5F). To understand the distribution of estimates and true sniff rate values, we constructed a 2D histogram of estimates and true values at various window sizes (Figure 6). We noticed that pervasive low frequency power in the LFPs is responsible for the inferiority of the frequency-domain method, especially at longer windows (Figure 6,1B, 1C *top*). That is, where the LFPs power spectrum predicts low sniff rate the time domain method which relies on peak finding is more accurate. Taken together, these results suggest that spectral estimations used on either the LFPs or sniffing signal may be imprecise estimates of the physiologically relevant oscillations and peak finding is a superior method for estimating oscillation rate.

**Figure 4:**
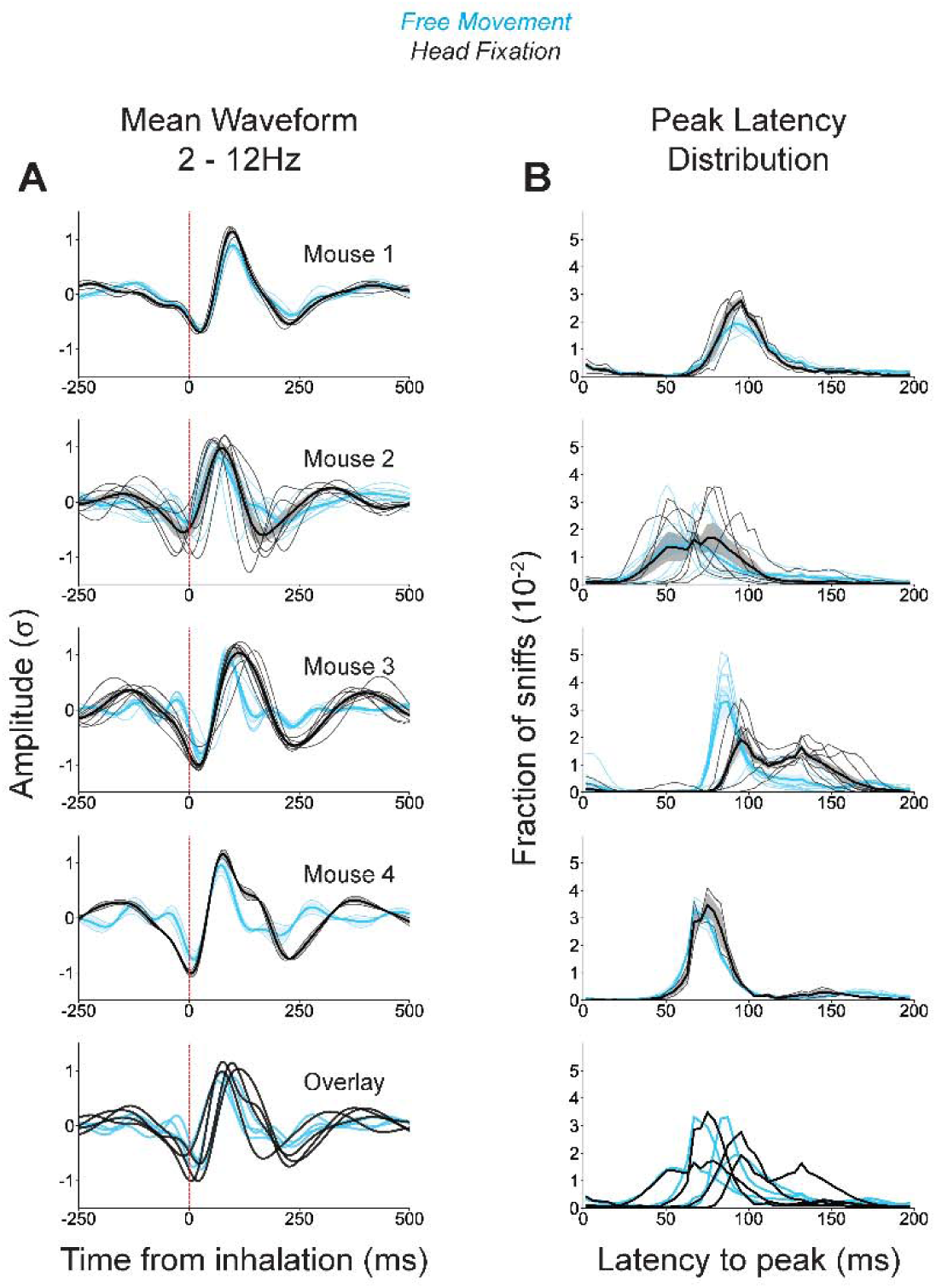
**A)** *Left.* Mean waveform of the unit variance scaled 2 – 12 Hz theta rhythms aligned to inhalation onset, taken from one session. *Right.* **B)** distribution of latencies to the peak in the LFPs following inhalation onset. Distributions were constructed on a sniff-by-sniff basis by finding the distance between inhalation onset and the first prominent LFP peak. **A, B)** Each of the four mice are shown with thin lines to represent the across-session within-mouse mean waveforms and latency distributions, the thicker line to show across-mouse grand mean, and shaded region is standard error of the grand mean computed across all the sessions. Overlay shows the mouse means. Head fixation represented in black with free movement in blue. Distributions differ significantly across mice (two-sample Kolmogorov Smirnov test; p < 0.01). In mouse 2 and 3, session distributions differ significantly (p < 0.01).

**Figure 5:**
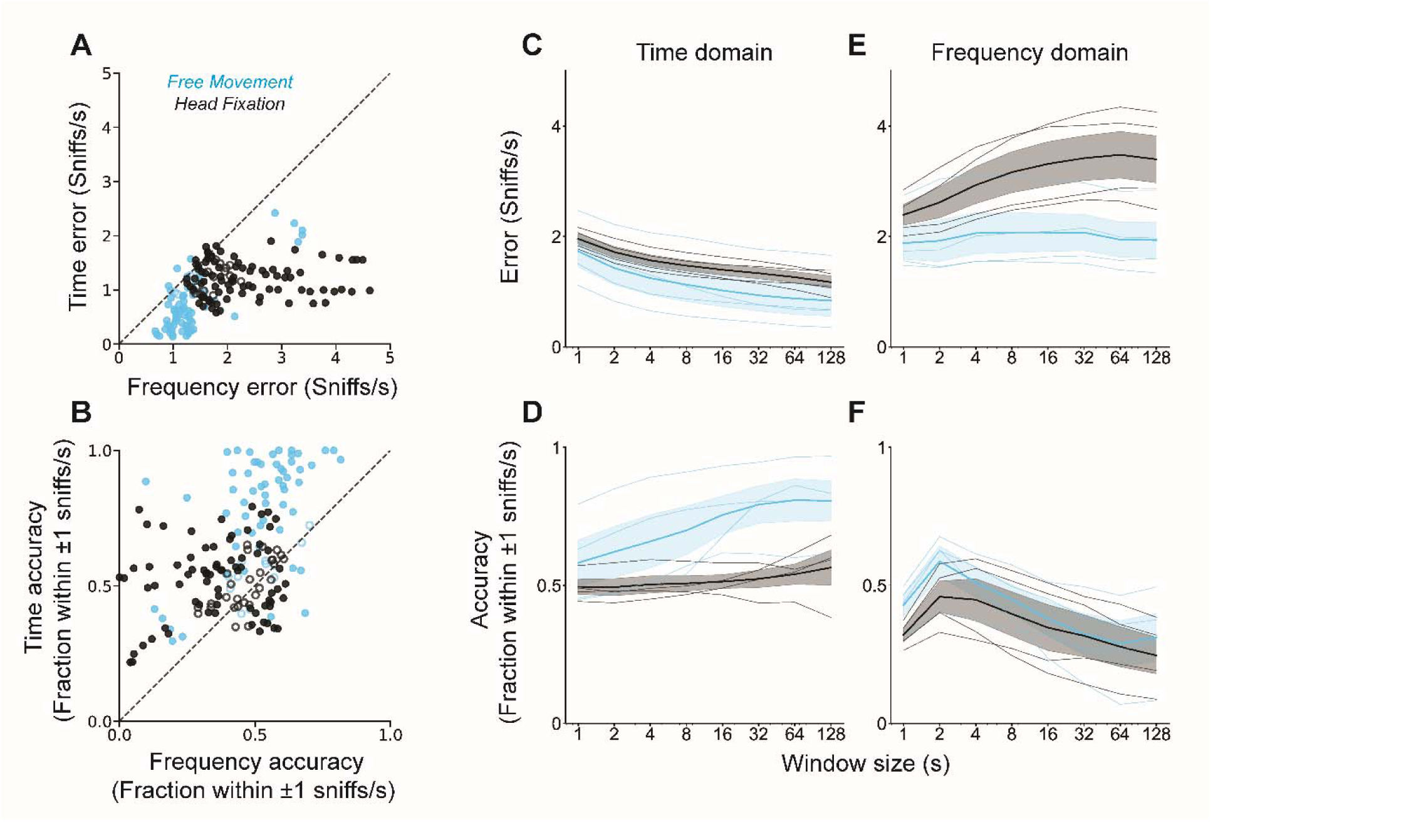
Respiratory rate can be recovered from LFPs. **A)** Scatterplot of the root mean squared error of the predicted respiratory rates using time and frequency domain methods. Filled dots represent sessions with significant deviation from equal errors across the two methods (p < 0.01, rank sum test), empty dots represent sessions where the error does not differ significantly between the two methods. **B)** Scatterplot of accuracy, as defined by the fraction of estimates within 1Hz of the true value, using time and frequency domain methods. **C)** Root mean squared error of the time domain estimates which find the mean reciprocal of interpeak intervals in the LFPs in increasing window sizes from 1 to 128 seconds. Thin lines represent mouse means with the thicker line for grand mean and shaded standard error of the mean. Head fixation in black, free movement in blue. **D)** Accuracy of the time domain estimates. **E)** Root mean squared error of the frequency domain estimates which estimates the frequency with maximum power between 2 and 12Hz in the LFPs in increasing window sizes. **F)** Accuracy of the frequency domain estimates.

**Figure 6:**
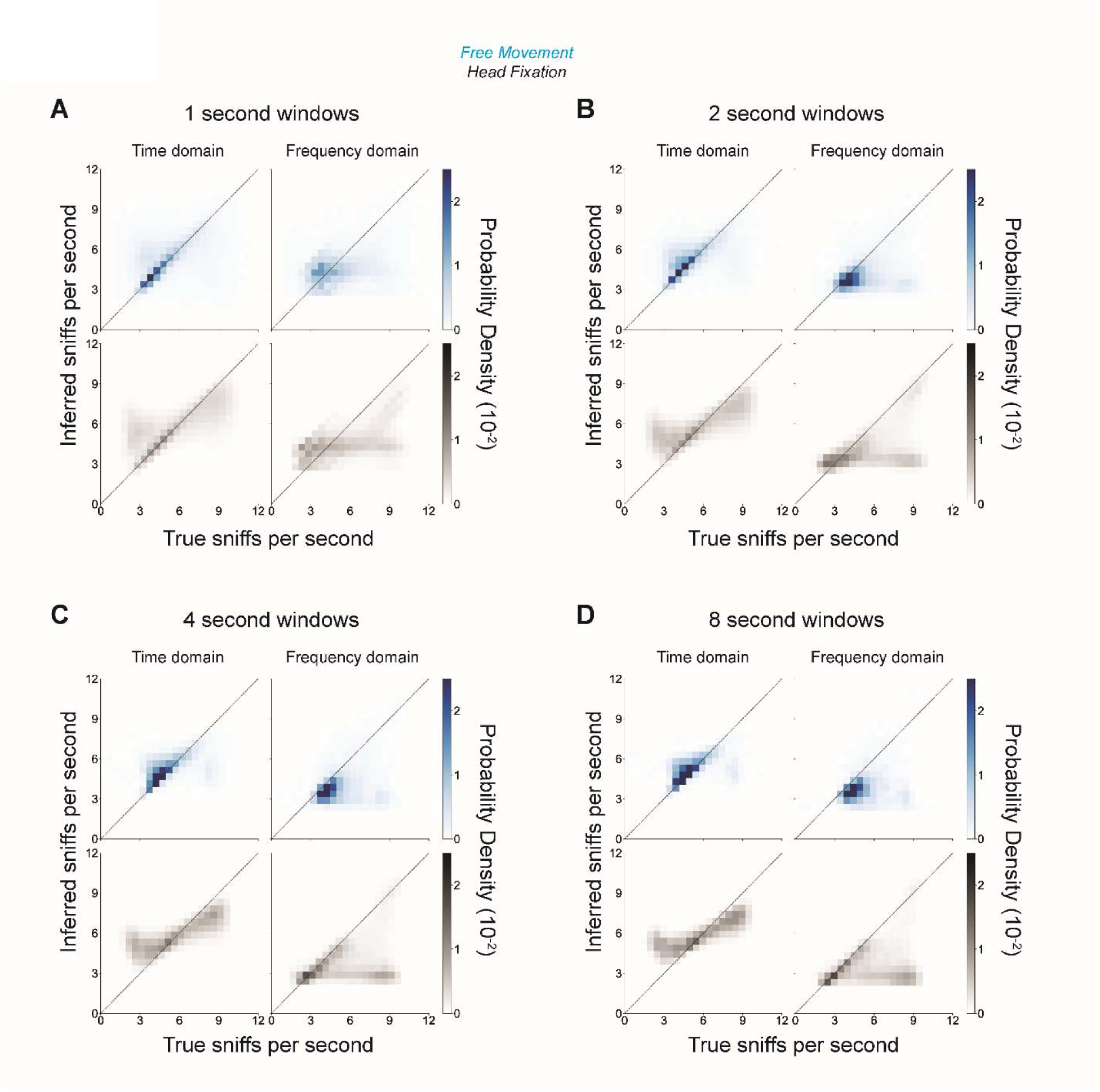
Probability density of the inferred vs true sniff rate shows the structure in the model errors. **A, B, C, D)** Heatmap shows the joint probability density of inferred vs true respiratory rates in 1, 2, 4 and 8 second sliding windows, respectively. Colormaps are scaled between 0 and the 95^th^ percentile of the density. Probability density of estimates in head fixation are shown ranging from white to black whereas in free movement density is shown ranging from white to blue.

## Discussion

Sniffing orchestrates odor dynamics and organizes the temporal structure of neural activity in the OB and elsewhere (Buonviso et al., 2006; Carey et al., 2009; Chaput et al., 1992; Heck et al., 2022; Karalis & Sirota, 2022; Kleinfeld et al., 2016; Macrides et al., 1982; Mori & Sakano, 2022; Pager, 1985; Tort et al., 2018). Here we asked whether OB LFPs can be used to infer the sniff signal. Despite the faithful coupling of sniffing and OB LFPs, inferring precise respiratory dynamics from OB LFPs presents several challenges. Our study demonstrated robust spectral coherence between OB LFPs and respiration, particularly in the theta band, which has previously been shown to track the respiratory cycle (Kay et al., 2009; Rojas-Líbano et al., 2014). We extended this finding into the time-domain where we describe the sniff-by-sniff variability in theta band rhythmic events—peaks and cycle lengths—with respect to inhalation onset. Furthermore, we show a modest respiratory alignment of beta band rhythms, which were disrupted upon averaging waveforms within sniff frequency bins. This result was not surprising considering the alignment of these rhythms to respiration were not robust across or within sessions. Further, the beta amplitude varies across behavioral states and is expressed differentially during context learning (Fourcaud-Trocmé et al., 2019). Finally, gamma oscillations are thought to play a critical role in the processing and integration of olfactory information, particularly under high task demands (Beshel et al., 2007; Losacco et al., 2020). We found that the amplitude envelope of these gamma oscillations aligns to inhalation onset, suggesting an inhalation aligned burst in gamma activity. Strikingly, the linear correlation between gamma bust length and instantaneous sniffing frequency was more robust in head fixation compared to free movement. However, the modest correlation in free movement rendered this frequency band insufficient for recovering respiratory frequency using our methods. Taken together, we used the theta band rhythms to attempt to infer respiratory dynamics. However, recovering the exact timing of inhalation onset and sniff frequency based on LFPs proved difficult due to variability in the latency of LFP peaks and inconsistencies across sniff cycles and individual animals.

One of the primary challenges arises from the inherent variability in the LFPs across respiratory cycles. While theta band rhythms often align with sniffing, containing a single peak per respiratory cycle, this is not always the case. This variability complicates the inference of precise respiratory frequency recovery on a cycle-by-cycle basis. Furthermore, our results show that inhalation aligned, sniff frequency binned and averaged theta epochs and gamma envelopes reliably contain a single peak per respiratory cycle, but the timing of these peaks relative to inhalation onset varies across individual sniffs and mice. This suggests that while LFPs may broadly represent respiratory rhythms, the exact timing of respiratory events such as inhalation onset may be masked by inter-session and inter-subject variability. This variability is likely due to different electrode locations in the OB across mice, and the advancing of the probes between sessions. However, even within a single session, the monotonic relationship between sniff duration and latency suggests that recovering the precise timing of inhalation is not as simple as subtracting a mean latency.

Moreover, we observed differences in LFP-respiratory relationships between free-moving and head-fixed conditions. In general, free movement resulted in better performance of both time and frequency domain methods in predicting respiratory frequency. This suggests that the animal’s behavioral state influences the quality of respiratory representation in LFPs. However, the gamma envelope and respiratory cycle length correlation was stronger in head-fixed compared to free movement. Free movement likely introduces additional physiological variables, such as increased respiratory modulation or altered neural dynamics, which may differentially enhance or diminish the coupling between LFPs across multiple frequency bands and respiration.

Despite these challenges, our findings suggest that OB LFPs can still serve as useful proxies for respiratory frequency, especially when averaged across multiple sniff cycles. By using time-domain methods to detect peak-to-peak intervals in LFPs, we were able to infer sniff frequency with a relatively low error margin during free movement. However, the variability in LFP-respiratory delay renders the recovery of precise inhalation events more challenging. This difficulty underscores the limitations of LFPs in capturing fine-grained temporal dynamics of respiration and highlights the need for complementary methods, such as thermistor-based sniff measurements, to accurately track respiratory events.

Future work should focus on refining models to account for the variability in LFP-respiratory coupling and explore ways to combine LFP-based methods with other non-invasive measures of respiration. Additionally, investigating the underlying neural mechanisms that contribute to the variability in LFP-respiratory alignment across different frequency bands and behavioral conditions may provide deeper insights into the relationship between respiration and OB neural activity.

In conclusion, while OB LFPs reliably track respiratory frequency, inferring precise inhalation timing remains a challenge due to the variability in LFP-respiratory delay. The use of time-domain methods shows promise in estimating respiratory frequency, particularly in free-moving conditions, but future work is needed to improve the temporal resolution of respiratory event detection from LFPs.

## ACKNOWLEDGEMENTS

We thank James Murray and Avinash Singh Balla for reviewing the manuscript

## AUTHOR”S CONTRIBUTIONS

Conceptualization, MS, SR, NH; Methodology, TF; Formal Analysis, SR; Data Collection, TF; Resources, MS; Data Curation, SR; Writing—Original Draft, SR; Writing—Review & Editing, SR, MS, NH Funding Acquisition, MS

## DATA AVAILABILITY

Data are archived at https://dandiarchive.org/dandiset/001433. All code and data for analysis are available at https://github.com/Sid-Rafilson-1617/sniff-LFP

